# An approach for accelerated isolation of genetically manipulated cell clones with reduced clonal variability

**DOI:** 10.1101/508721

**Authors:** Natania Casden, Oded Behar

## Abstract

Genomic editing methods, such as the CRISPR/Cas9 system, are routinely used to study gene function in somatic cells. Due to the heterogeneity of mutations, it is necessary to purify cell clones grown in high dilution until colony formation, which can be a time consuming process. Here we tested a modified approach in which we seeded cells in high dilution, together with non-edited carrier cells. As a comparison, cells were also grown in high dilution with conditioned medium from a high density culture. When using carrier cells or conditioned medium, the formation of cell colonies is accelerated. Additionally, clones grown with carrier cells are more similar to the parental lines in terms of their tumorigenic properties. Surprisingly, key signaling cascades are very divergent between clones isolated from low density cultures, even with conditioned medium, in contrast to clones isolated with carrier cells. Thus, our study uncovers a significant limitation using the common approach of isolating cell clones following genetic modifications and suggests an alternative method that mitigates the problem of heterogeneity of gene expression between clones.

**Summary statement:** We show here that standard methods of generating cell clones after CRISPR/Cas9 treatment result in highly divergent signaling cascades among clones. We demonstrate here an improved alternative approach.

## Introduction

In recent years, a number of genomic editing methods have been developed which enable modification of the genome in somatic cells. The most recent and widely used genome-editing technology employs the CRISPR/Cas9 system (Makarova et al., 2011). This system employs a single guide (sg) RNA which targets a specific genomic DNA sequence and also serves as a scaffold domain which binds the Cas9 nuclease (Makarova et al., 2011). This method has been used to generate targeted mutations in numerous model organisms such as zebrafish, mouse, rat, and human (Fogarty et al., 2017; Li et al., 2016; Qin et al., 2011; Shao et al., 2014). It is also used to generate somatic mutations in cells in order to test specific gene function (Munoz et al., 2014; Shah et al., 2016). Mutations induced by the CRISPR/Cas9 system can result in diverse sequences in each cell in a given population and even in the same cell in each allele. Therefore, it is imperative to isolate individual cell colonies in order to obtain a pure sub-line of the original parental cells containing identical mutations. Common methods of acquiring pure sub-lines are plating the genetically manipulated cell population in low density so as to obtain individual colonies or alternatively, by FACS sorting them into 96 well plates so that each well receives only one cell. In both methods, a uniform sub-line will be isolated and sequenced in order to verify the exact nature of the mutation (Bauer et al., 2015). The difficulty with this approach is the stage in which cells are grown in low density or in complete isolation. When grown in low density, most cells require a long period of time to divide and form colonies. In addition, the ability of cells to grow in effective isolation may apply selection pressure toward subpopulations of cells that are more suited to grow under such conditions, one of the characteristics of more tumorigenic cells. We have therefore decided to test the effects of this selective pressure on cell clones obtained and the possibility that the addition of non-modified carrier cells of the same line might reduce selection and accelerate the growth of individual cells after plating.

## Results

### Co-plating with carrier cells increases the percentage of colony formation and accelerates their formation

In order to analyze the effects of genetic manipulation on specific cells, it is critical to isolate homogenous cell populations. It is customary to acquire these homogenous populations by seeding the CRISPR treated cells in high dilution. Since most cells do not grow well in isolation, only a small number of cells of the original pool are able to thrive and form colonies. We hypothesized that culturing carrier cells-cells of the same line but with no antibiotic resistance, in moderate density together with the highly diluted, antibiotic resistant modified cells might accelerate colony formation and increase the percentage of cells forming colonies. In order to test this idea, we generated a pool of puromycin resistant U87MG, U251, and NIH3T3 cells. We then seeded these cells in high dilution in the presence of a large number of non-resistant parental cells. When the cultures reached high density, we added puromycin as a selection medium to eliminate the non-resistant carrier cells. In parallel, we plated the antibiotic resistant U87MG, U251, and NIH3T3 cells in high dilution without carrier cells. In all three lines, there was an acceleration of colony formation (Fig 1A, B, C, and D) of the puromycin resistant cells grown in the presence of carrier cells. In addition, the number of individual cells able to form cell colonies in the presence of carrier cells was dramatically increased in each line tested (Fig 1 E, F, and G) in comparison to the cells plated without carrier cells in a low density culture. To ensure that our cell clones were not contaminated with carrier cells, we used a cell pool of U251 cells infected with a lentivirus containing the mCherry red fluorescent gene as carriers and seeded 400 puromycin resistant U251 cells. We then repeated the process of clone isolation. Almost all cells were mCherry positive at the time of plating, however there was no fluorescence identified in the isolated, expanded clones (Fig S1).

**Figure 1:**
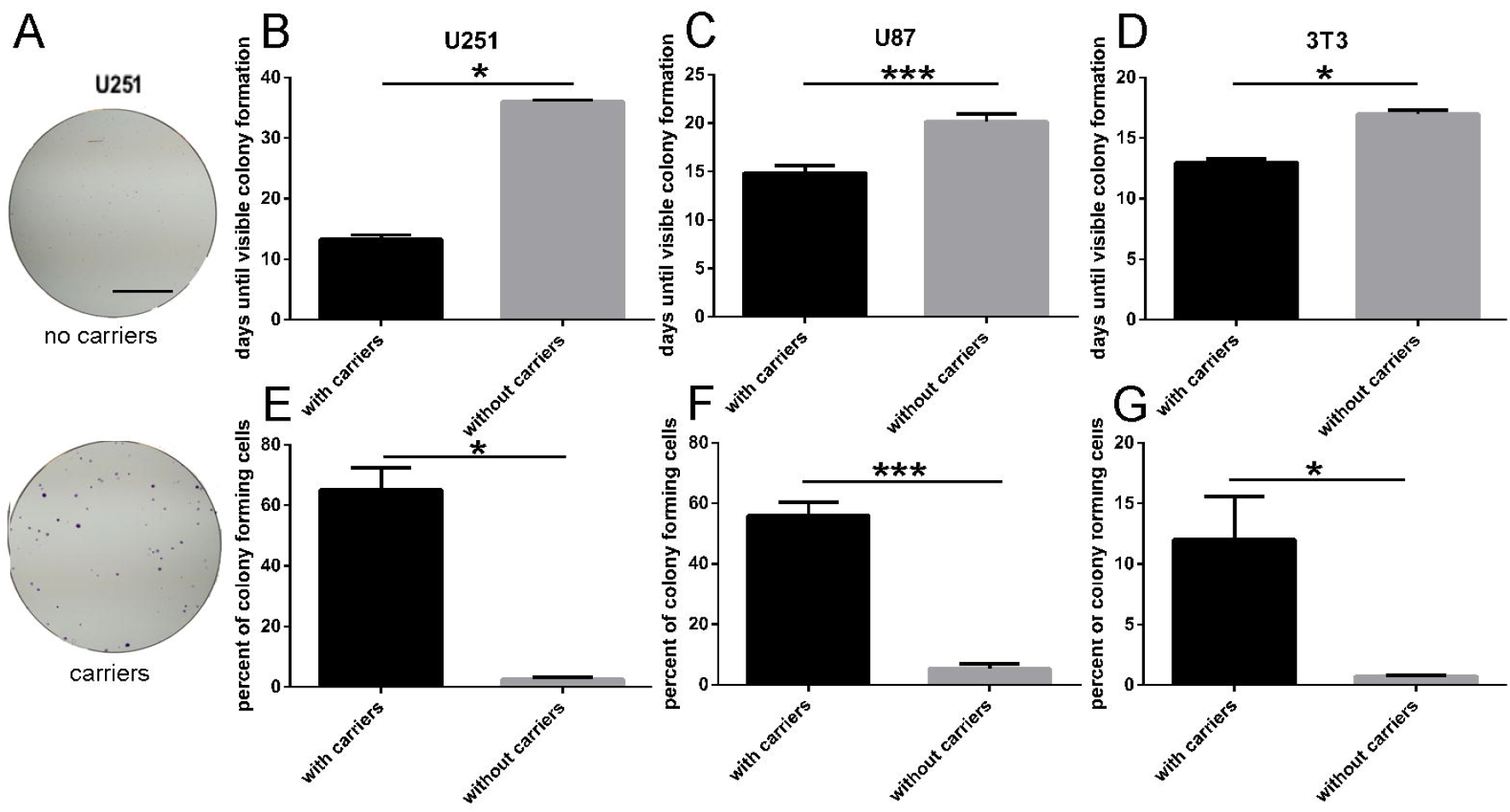
increased number of colonies in accelerated rate when cells are seeded with carriers. (A-G) Colony formation assay experiments were performed for U251 (n=3 (without carriers), 4 (with carriers)), U87MG (n=5 (without carriers), 8 (with carriers)) and NIH3T3 (n=3 (without carriers), 4 (with carriers)). A) representative example results. In upper panel 400 cells were seeded without, and lower panel 100 cells were seeded with carriers. The plates were stained with crystal violet after 14 days. (B, C, D) Colonies were counted for each well; data represent the median of experiments (the p values were calculated with the one-tailed Wilcoxon-Mann-Whitney Test). (E, F, G) days until colonies were of sufficient size to lift are presented. Data represents the median of experiments (the p values were calculated with the one-tailed Wilcoxon-Mann-Whitney Test). Scale bar 2cm.

### Co-plating with carrier cells reduces the frequency of cell senescence

What can be the explanation of the difference in the ability to form cell clones with and without carriers? One possibility is the entry into cell senescence. To test if this is indeed the case among cells seeded in low density, we seeded highly diluted U87MG cells with and without carrier cells. We then tested senescence by staining for senescence-associated beta-galactosidase (SA-β-gal). The percentage of SA-β-gal positive cells was much higher in individual cells and small cell colonies (2-8 cells) in cultures grown with no carrier cells than in those grown with carrier cells, after the selection process (Fig 2). This indicates that seeding cells with carriers helps to prevent the modified cells from entering into senescence and therefore enables more cells in the given population to generate viable colonies.

**Figure 2:**
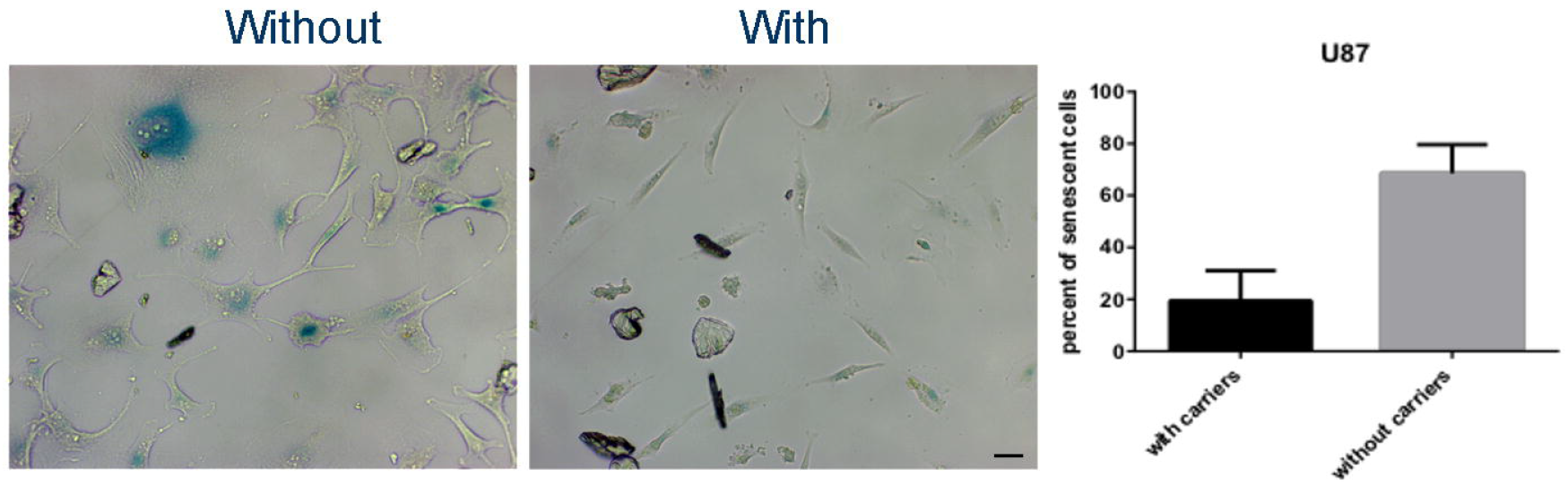
Reduced percentile of U87MG cells entering senescence when seeded with carriers. U87MG cells were seeded in low density or with carriers. After 14 days cells were stained for sA-β-gal assay. The percentile of colonies with β-gal positive cells is presented. Scale bar 200μM.

### Clones isolated without carrier cells perform much better in clonogenic assays

Thus far we have shown that when plating U87MG, U251, and NIH3T3 cells in high dilution, only a small fraction of them form cell colonies. We further showed that by using carrier cells, more cells of the initial population are able to form clones. However, is there a difference between the two types of cell clones achieved by the different methods? The colony formation assay is one of the assays used to test tumorigenic potential of cells. We decided to test whether clones isolated by the two different methods have the same ability to form cell clones if re-cultured a second time in low density with no carrier cells. We isolated U87MG and U251 clones from cultures grown with and without carriers, and recultured them by seeding them in low density with no carrier cells. We then counted the number of colonies formed from the individual cells (Fig 3A, B, and C). In both the U87MG and U251 lines, a dramatically higher number of colonies was obtained from clones that had originally been isolated from cultures grown without carriers. This indicates that the process of seeding cells in effective isolation leads to the formation of more clonogenic colonies when recultured in high dilution than cells grown with carrier cells, which showed a more limited ability to form new colonies in low density cultures.

**Figure 3:**
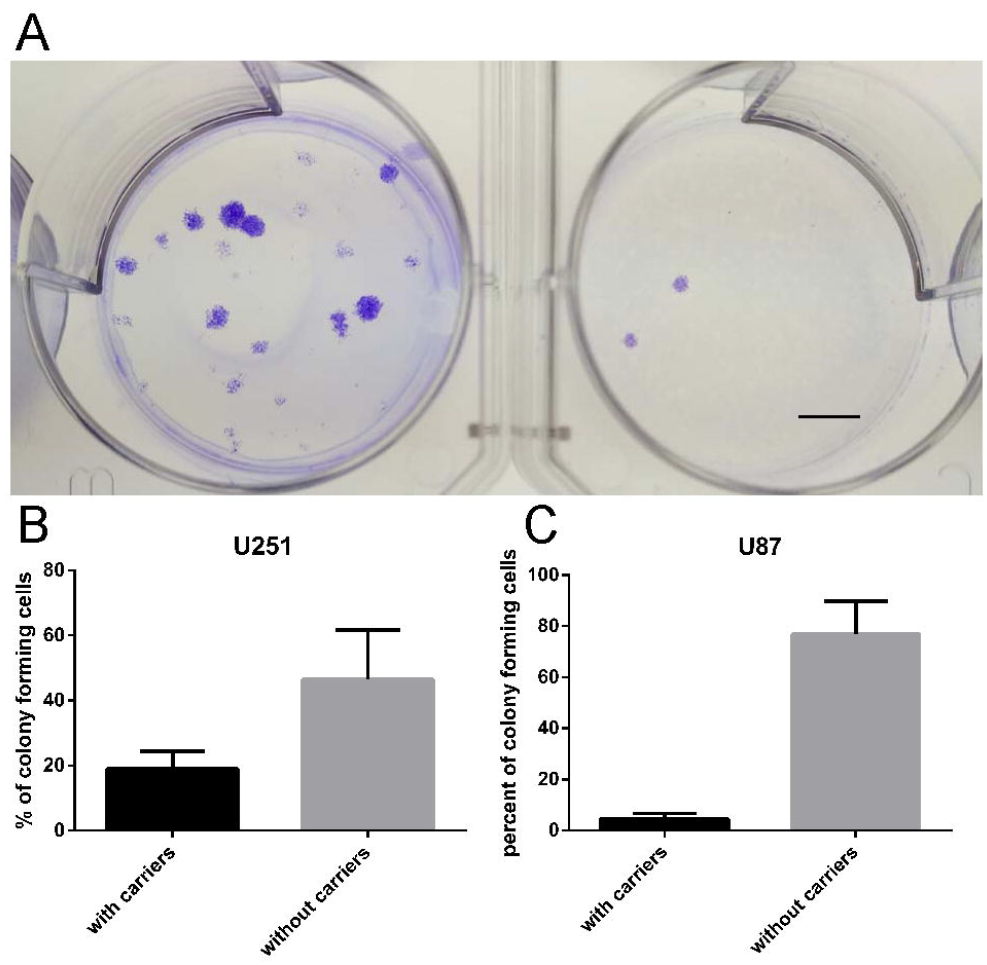
Colony formation assay of clones isolated with or without carriers. A, B, C) Colony formation assay experiments with were performed in triplicate for clones isolated with or without carriers. A) a representative example of U251 clone isolated without (left) or with carriers. Average clonal formation of U87MG (n=12 (without carriers), 10 (with carriers) (B) and U251 (n=3 (without carriers), 4 (with carriers)) (C) are shown. Scale bar 0.5cm.

### Clones isolated with carrier cells can be used in CRISPR/Cas9 experiments

We have recently demonstrated that CRISPR/Cas9 targeted mutations in the signal sequence of Sema4B in different glioma lines had no effect on cell proliferation or ability to form clones (Peretz et al., 2018). We used the same gRNA to generate mutations in the signal sequence of Sema4B and used our carrier approach to isolate cell clones in order to show that this method can be used in conjunction with CRISPR/cas9 editing. In addition to clones grown with and without carriers, we also isolated clones grown in conditioned medium collected from dense cultures. Both conditioned medium and the use of carriers accelerated clonal isolation and resulted in a higher percentage of clone formation (Fig 4 A, B). Similar results were also obtained when using U87MG (Fig S2). A few of these clones isolated with carriers were analyzed, sequence verified, and tested for cell proliferation and clonability and were presented in a separated study (Peretz et al., 2018). We then tested the individual clones isolated by the different methods in clonogenic assays. As before, cell clones isolated with carrier cells were able to form clones similar to parental line. In contrast, clones isolated without carriers were more effective in the clonogenic assays, indicating that these clones are more tumorigenic (Fig 4C). Clones isolated in conditioned medium shown the same trend as clones isolated without carriers, however the difference between them and clones isolated with carriers did not reach statistical significance.

**Figure 4:**
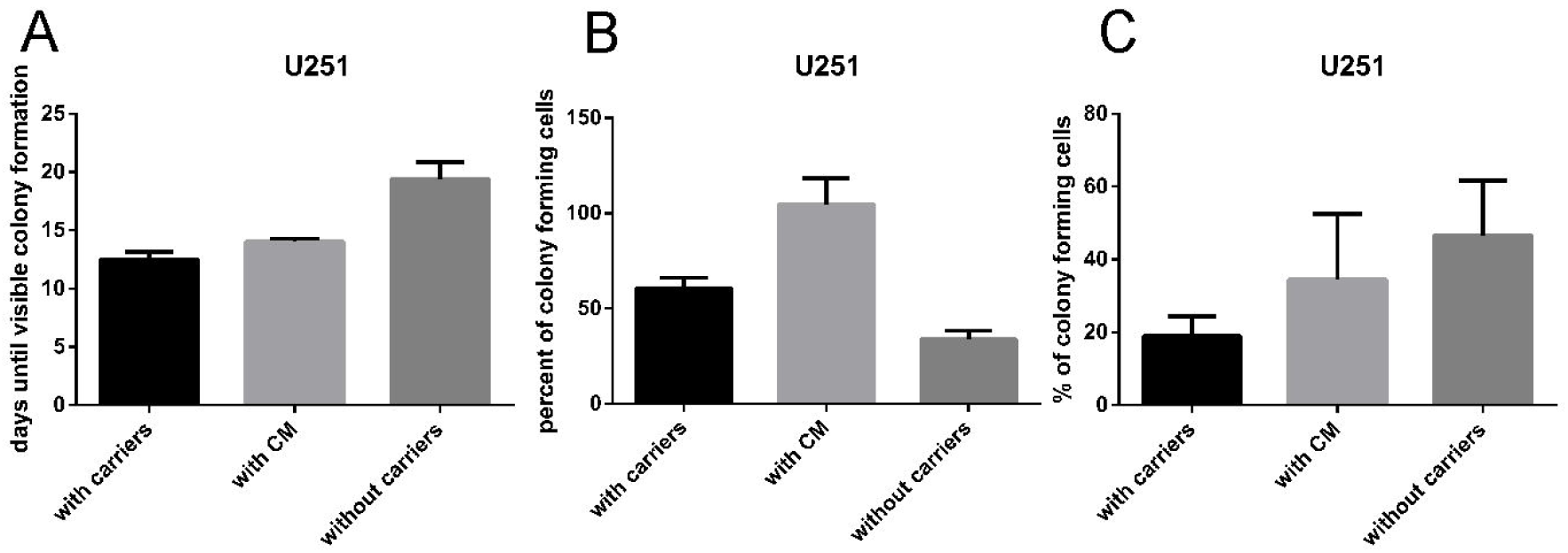
Isolation of cell clones after CRISPR/Cas9 mutation in the Sema4B signal sequence. U251 cells were treated with lentiviral particles coding for Cas9 and gRNA targeting the signal sequence of Sema4B. In upper panel, cells were seeded with carrier cells (n=6), with conditioned medium of parental cells (n=3), or alone (n=6). The time until colonies were ready to be isolated is shown in A. The percentile of cells forming colonies is shown in B. From each set of conditions we selected 3 clones and tested their clonogenic potential C. Although on average, cell clones isolated from conditioned medium give more colonies than clones isolated with carriers, the difference was not statistically significant.

### Signaling cascades in colonies isolated by co-plating are more uniform and more similar to the parental lines

When we apply genomic editing and isolate clones, it is generally assumed that clones with the same genetic modification will be similar to each other. Thus, when comparing genetically modified clones to control lines, any disparity in signaling cascades will be attributed to the genome manipulation. Since we have seen a significant difference between clones isolated with and without carriers with respect to their tumorigenic potential, we questioned whether we would find a significant variation in major signaling cascades between the different clones isolated by the two methods. We chose to test the activation state of major MAPK signaling proteins including MAPK44/42, p38 MAPK, and JNK in our parental cell pool and in the individual clones isolated with or without carriers (Fig 5, Fig S3). Overall, clones isolated without carriers tend to be more divergent from one another and less similar to the parental line. In contrast, cells isolated with carriers are less divergent from one another and more similar to the parental line.

**Figure 5:**
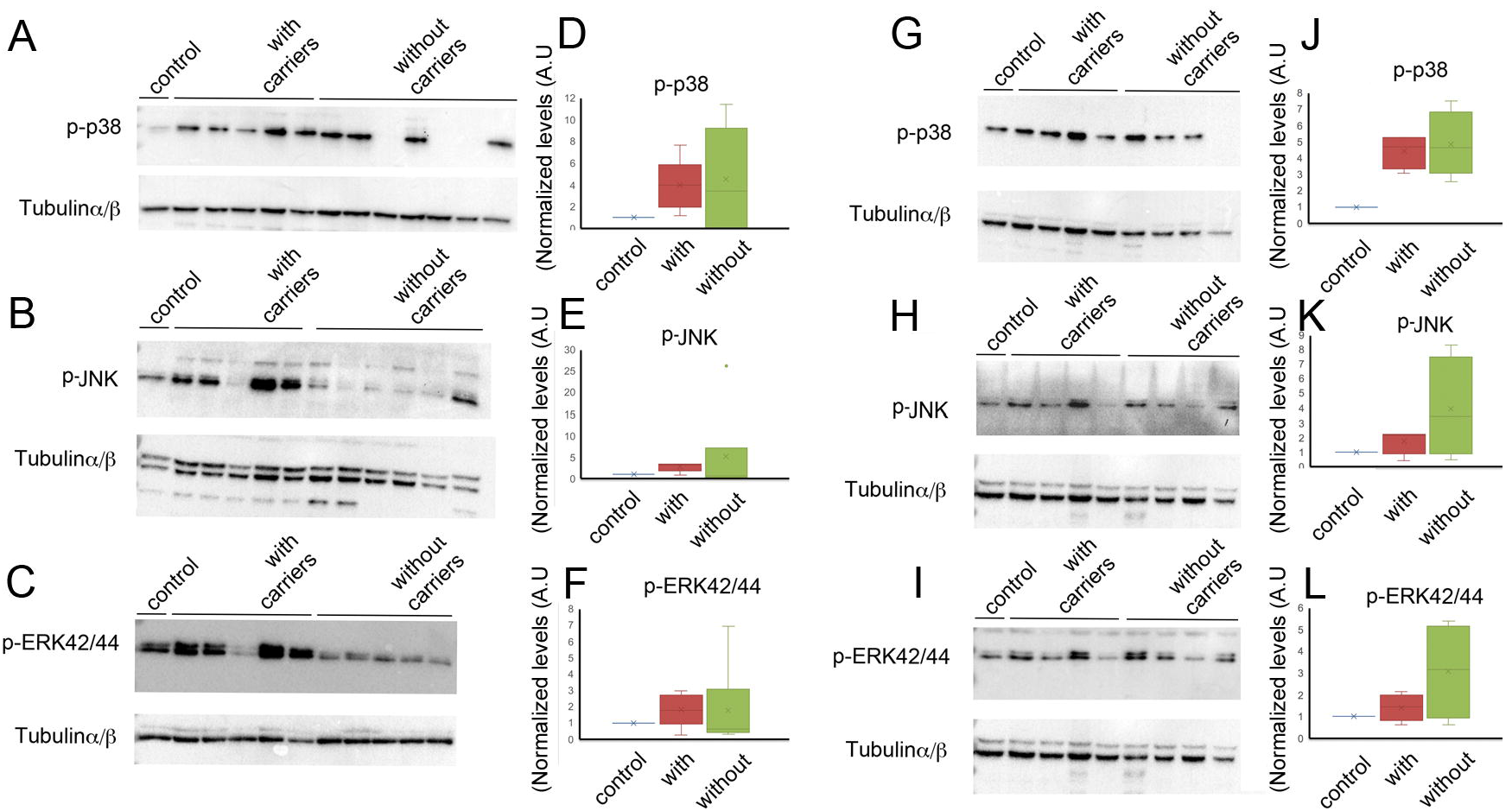
MAPK signaling cascades are altered more dramatically in clones isolated without carriers. We isolated and tested 5 U87MG clones with and 8 clones without carriers (representative western blots are shown in A). To compare the normalized (relative to Tubulin) intensity of signals we calculated the median of 3-4 blots for each clone (3-4 different protein extractions for each clone). The chart shows distribution of data into quartiles, highlighting the mean and outliers. The lines above and below the quartiles indicate variability outside the upper and lower quartiles (B). We also isolated and tested 4 U251 clones with and 4 without carriers (representative western blots are shown in C). To compare the normalized (relative to Tubulin) intensity of signals we calculated the median of 4 blots for each clone (4 different protein extractions for each clone). The distribution of data and mean of all clones with or without carriers for each cell clone is presented (D).

To get more quantitative information regarding the possible changes in U87MG cells grown with or without carries, we selected 24 genes which are associated with tumorigenicity. Out of these genes, 5 genes were not expressed in U87MG and 8 were similar in all conditions. However, gene was expressed more highly overall, and 10 genes were more divergent between clones grown without carriers (Fig 6A, S4). To further validate these results, we selected 8 of the genes found to be highly variable in U87MG clones, and used them to test U251 clones. This time we tested cell clones isolated with conditioned medium as well (Fig 6B, S5). In this case, 6 out of the 8 genes were much more divergent in clones isolated without carriers, and importantly, in cell clones grown in conditioned medium as well.

**Figure 6:**
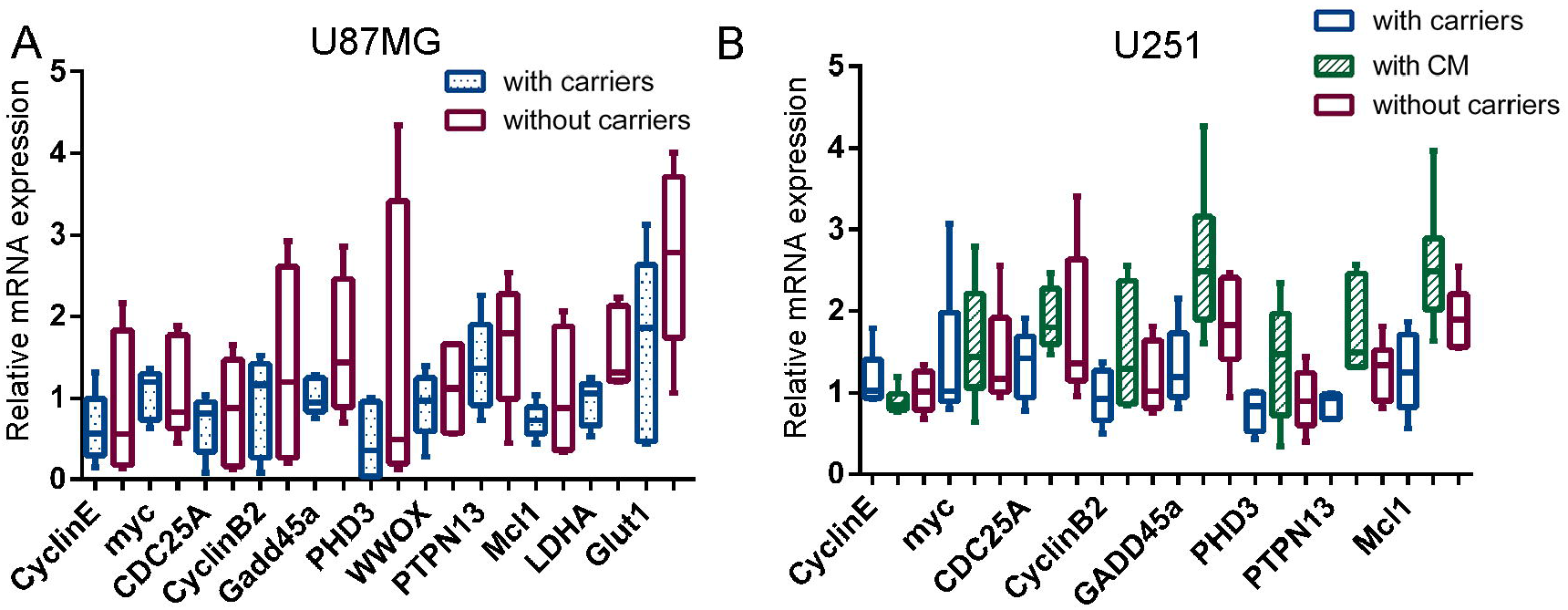
testing cell clone heterogeneity at the RNA expression level. qPCR analysis of a set of cancer associated genes in U87MG (A) and U251 cell clones (B). Note the reduced variability of gene expression detected in clones isolated with carriers as compared to cell clones isolated without carriers or with conditioned medium.

## Discussion

Genomic editing methods such as CRISPR/Cas9 system may be used to generate genomic mutations, insertions, and deletions in a targeted manner. The events that underlie these mutations, deletions, and insertions requiring local repair, result in a series of alleles with a variety of indels at the target sites. Thus, isolation of cell clones to obtain genetically homogenous sub-lines is necessary. The current method of achieving this, low-density cell plating, is necessary in order to obtain isolated colonies. Once genomic manipulation in each clone is validated, it is generally assumed that each of the clones are identical to one another. Here we show that when growing cells in high dilution, the sub-lines generated display more tumorigenic characteristics than do the parental lines. Moreover, major signaling transduction cascades such as p38, JNK, and ERK42/44, as well as important cell cycle, metabolism, and stress related genes also show increased variability among clones. Thus, comparing these sub-lines to the parental line or other sub-lines with different genomic editing targets is potentially misleading. Importantly, although growing cells in conditioned medium leads to accelerated formation of cell clones, the variability of gene expression in these clones is similar to, or even higher than clones isolated without carriers. These limitations are mitigated when using our approach in which cells are plated with carriers, and the sublines generated are more similar to the parental lines and to each other. This conclusion is supported both by clonogenic assays of cells grown with carriers, which display more similarity to the parental lines, and the signaling cascades and cancer associated genes tested which are less variable among clones grown with carriers and more closely resemble the parental lines. Thus, we propose that our strategy makes it possible to isolate modified cell colonies more quickly, and reduces the selection pressure on these cells that generally leads to clones with more tumorigenic properties.

## Materials and methods

Antibodies: Antibodies specific for phospho-SAPK/JNK (Thr183/Tyr185) (#9251, dilution 1:1000), phospho-MAPK42/44 (Threonine 202 and Tyrosine 202 of human MAP kinase) (#4376, dilution 1:1000), Phospho-p38 MAPK (Thr180/Tyr182) (#4511, dilution 1:1000), and α/β-Tubulin (#2148, dilution 1:1000) were purchased from Cell Signaling Technology (Beverly, MA, USA).

### Cell Culture

Human glioblastoma cell lines U87MG, U251, and NIH3T3 are from the ATCC (Virginia, USA) and were grown in medium according to their guidelines.

#### Sub-line isolation using co-plating with carriers

U87MG, U251, and NIH3T3 were infected with a pLKO1 lentivirus, using one infecting unit/cell and selected with puromycin to generate resistant cell pools for each line. To generate clones in high dilution, we seeded cells in 10cm plates as such: U87MG- 3 plates of 800 cells each, and 2 plates of 500 cells each; U251- 3 plates of 400 cells each; NIH3T3- 3 plates of 2,000 cells each). To generate clones with carriers, we plated cells with parental carrier cells in 10cm plates as such: U87MG- 8 plates each of 100 cells with 100,000 carrier cells; U251- 4 plates each of 100 cells with 50,000 carrier cells; NIH3T3- 4 plates each of 100 cells with 150,000 carrier cells. Once the cells with carriers reached high density we eliminated the carrier cells with one round of puromycin selection (U87MG- 4μg/ml, U251- 2μg/ml, NIH3T3- 1μg/ml) for 2 days. To generate U87MG clones in conditioned medium, we seeded 100 cells each in 6 10cm plates in medium taken from dense U87MG cultures and filtered through a 22micron strainer. To generate clones from U251 cells treated with CRISPR/cas9, we seeded cells in 10cm plates as such: 6 plates each of 200 cells alone, 3 plates each of 50 cells with conditioned medium, and 6 plates each of 50 cells with 50,000 carrier cells. Colonies were lifted with Trypsin C (Biological Industries #030531) and transferred to 96 well plates. Colonies lifted from cultures grown with carrier cells were transferred to wells containing puromycin to eliminate any remaining carrier cells.

#### Florescent labeled carrier cells

U251 cells were infected with an FUW based vector in which the coding region of hygro-cherry was cloned using an ECoRI site. The cells were selected with Hygromycin (200μg/ml) for a week to ensure that all cells express the vector. Cells were used as carrier cells in some control experiments.

### Colony formation assay

To test the ability of the first round of clones to form new clones, we seeded cells from each clone in 3cm plates in triplicate as such: U87MG cells: 2 sets of samples were seeded from 5 clones isolated from cultures with carriers- 1 set with 40 cells each, and 1 set with 25 cells each; 2 sets of samples were seeded from 6 clones isolated from cultures grown in low density- 1 set with 40 cells each, and 1 set with 25 cells each. U251 cells: 30 cells each were seeded from 4 clones grown with carriers and from 3 clones grown without carriers. U251 cells treated with CRISPR/cas9: 20 cells each were seeded from 3 clones grown with carriers, from 3 clones grown without carriers, and from 3 clones grown in conditioned medium. After incubation and colony formation, cells were fixed with crystal violet for 5 minutes and washed 3 times with PBS. The colonies were counted.

#### Senescence-associated beta-galactosidase (sA-β-gal) staining

U87MG cells were seeded in 4cm^2^ wells as such: 50 cells without carrier cells, and 10 cells with 6,500 carrier cells, were seeded in 4 wells each. After 2 weeks, cells were fixed with 0.2% glutaraldehyde in PBS at room temperature for 10 minutes, and washed twice with PBS and once with 1mM PBS/MgCl2 for 5 minutes each. Cells were incubated for two hours at 37°C in X-Gal solution (PBS-MgCl2 (1mM) pH 6, NaCl (1.5M), K3Fe (100mM), K4Fe (100mM), and X-Gal (40mM)) filtered with a 45micron strainer and incubated at 37°C before use. The cells were then washed 3 times in PBS for 5 minutes and colonies were counted.

### Immunoblots

Cells were harvested in RIPA lysis and extraction buffer (Thermo Fischer Scientific), 0.2mM sodium orthovanadate (Sigma), and protease inhibitor cocktail (25x, Sigma). Cells were collected with a cell scraper, then vortexed and incubated on ice for 5 minutes, 4 times. The lysate was then centrifuged at 20,000 x g for 15 min and the pellets discarded. Samples were boiled in 1x SDS sample buffer, separated by SDS-10% polyacrylamide gel electrophoresis (PAGE) and blotted onto PVDF membranes (Millipore). The membranes were incubated in 5% fat-free milk in TBST (10mM Tris-HCL pH 7.4, 150mM NaCl, 0.1% Tween 20) for 1h and then 5% BSA and 0.01% NaN3 in TBST containing various dilutions of primary antibodies for 18h at 4°C. The membranes were washed three times with TBST for 5 min each before and after incubation with secondary antibody. The proteins were detected with an appropriate secondary antibody (1h, RT) coupled with a horseradish peroxidase-conjugated goat anti-rabbit antibody and visualized by chemiluminescence according to the manufacturer’s instructions (West Pico, Pierce). Signal strength of protein tested for each clone was normalized to αβ-tubulin levels and compared to the parental line using the GelQuant software program by BioChem Lab Solutions.

### CRISPR-Cas9

Templates for gRNA were cloned into lentiCRISPR v2 (Addgene Plasmid #52961). For this experiment we used a gRNA sequence targeting the first coding exon downstream to the first ATG start site: The sequences are:

Sema4B gRNA1:
5’-CGCACCGCGATGGGCCTG
5’-CAGGCCCATCGCGGTGCGC

The Cas9-gRNA plasmid was used to generate lentiviral particles. Lentiviral titer was estimate as follows: A day after infection, puromycin selection was applied and a day after selection the number of cells in each well was counted. From these results we estimated the transducing units/ml for each set of gRNA prep. In these experiments we used an estimate of one infection unit/cell.

### qPCR analysis

RNA was extracted using Quick-RNA MiniPrep (ZYMO R1055). Total RNA (1μg, as determined by Nanodrop, purity 260/280 above 1.9) in a total volume of 20μL was reverse-transcribed with the qScript cDNA Synthesis Kit (Quantabio, Beverly, MA) according to the manufacturer’s instructions. The resulting cDNA reaction mix was then diluted 4-8x in double-distilled water. Real-time quantitative PCR (qPCR) was performed with the SYBR Green mix (Roche) according to the manufacturer’s instructions. The specific primers were as follows:

BCL2: TTGCTTTACGTGGCCTGTTTC and GAAGACCCTGAAGGACAGCCAT
BCLW: GACCCGTGAGATCCCTAACCT and TGGGGCCTTTCATCCTCCT
BCLXL: ACCTGAATGACCACCTAGAG and GCTGCATTGTTCCCATAG
CDC25A: CCCT ACCTCAGAAGCTGTTGGGA and GAGTGCAGGCAGCCACGAGA
CyclinB2: AAGCTCAGAACACCAAAGTTCC and TGTCCTCGATTTTGCAGAGCA
CyclinE: ATCAGCACTTTCTTGAGCAAC and TTGTGCCAAGTAAAAGGTCTC
Gadd45a: GAGAGCAGAAGACCGAAAGGA and CAGTGATCGTGCGCTGACT
LDHA: TTGACCTACGTGGCTTGGAAG and GGTAACGGAATCGGGCTGAAT
Mcl1: TAGTTAAACAAAGAGGCTGG and ATAAACTGGTTTTGGTGGTG
myc: CTGGTGCTCCATGAGGAGA and TCCAGCAGAAGGTGATCCA
p21: CTGGAGACTCTC AGGGTCGAA and GGCGTTTGGAGTGGTAGAAATCT
p53: GAGTATTTGGATGACAGAAACACTTT and CCAGTGTGATGATGGTGAGG
PHD3: TCCTGCGGATATTTCCAGAGG and GGTTCCTACGATCTGACCAGAA
PTPN13: CAAGGAARGACCTTGGAGGA and CAGCAGACCTCTTTGAGCTG
sestrin2: AAGGACTACCTGCGGTTCG and CGCCCAGAGGACATCAGTG
WISP: CACGCTCCTATCAACCCAAGT and GCACCTATTGTCCATGCAAACTC
WWOX: TCCTCAGAGTCCCATCGATTT and TTTTGTTGGAGAGAGGCGAC
SMAD5 TCTCCAAACAGCCCTTATCCC and GCAGGAGGAGGCGTATCAG
Glut1 TCTGGCATCAACGCTGCTTTC and CGATACCGGAGCCAATGGT
SMAD3 TGGACTTAGGAGACGGCAGTCC and CTTCTGAGACCCTCCTGAGTAGG
TCF-7/TCF1 AGCTTTCTCCACTCTACGAACA and AATCCAGAGAGATCGGGGGTC
ERG: CACGAACGAGCGCAGAGTTAT and CTGTACTCCATAGCGTAGGATCT
MAPK14: TGACACAAAAACGGGGTTACG and GGTCTGGAGAGCTTCTTCACT
ERbB4: TGCCCTACAGAGCCCCAACTA and GCTTGCGTAGGGTGCCATTAC

The relative quantification of gene expression levels were normalized to GAPDH expression level using the following gene specific primers; 5’- ATGGGGAAGGTGAAGGTCGG -3’ and 5’ - TGACGGTGCCATGGAATTTG-3’ and the ΔΔCt method was applied (Schmittgen and Livak, 2008).

#### Statistical analysis

*Colony formation assay, Anchorage-independent growth assay* and SA-β-gal staining: Data are presented as means ± s.e.m. The p values were calculated with the Wilcoxon-Mann-Whitney Test after the data was confirmed as fulfilling the criteria. Symbols are as follows: *: P<0.05, **: P<0.001, ***: P<0.0001.

## Acknowledgment

We are grateful to Dr. Norman Grover (Department of Experimental Medicine, The Hebrew University) for helpful advice regarding the statistical analyses.

---

**Figure S1:**A) Culture of fluorescently labeled U251 carrier cells. B) Cell clone isolated using fluorescent expression carriers. Note that no carrier cells can be detected. Scale bar 1000μM.

**Figure S2:**Colony formation assay experiments were performed for U87MG cells with carriers, without carriers, and with conditioned medium of parental cells.

**Figure S3:**Cell clone variation is lower when using a carrier strategy. Western blots of 3 independent experiments testing U87MG cell clones isolated with or without carrier cells testing for p-p38. M- marker, P – starting cell pool, 1-5 with – clones isolated with carrier cells. 1-8 without – clones isolated without carrier cells. Among these clones – 2, 4, 6, 7 and 8 are shown in all blots.

**Figure S4: qPCR Analysis of additional genes which were used to test cell clone heterogeneity:**qPCR analysis of additional sets of cancer associated genes in U87MG. Heterogeneity is compared between cell clones isolated with or without carriers.

**Figure S5:**qPCR analysis of 6 U251 cell clones each isolated with carriers, with conditioned medium, or without carriers. Note the divergent expression between clones isolated with conditioned medium or without carriers and in each clone with respect to the different genes.

